# Benchmarking single-cell hashtag oligo demultiplexing methods

**DOI:** 10.1101/2022.12.20.521313

**Authors:** George Howitt, Yuzhou Feng, Lucas Tobar, Dane Vassiliadis, Peter Hickey, Mark A. Dawson, Sarath Ranganathan, Shivanthan Shanthikumar, Melanie Neeland, Jovana Maksimovic, Alicia Oshlack

**Affiliations:** Computational Biology Program, Peter MacCallum Cancer Centre, Parkville, VIC, Australia; Sir Peter MacCallum Department of Oncology, University of Melbourne, Parkville, VIC, Australia; Respiratory Diseases, Murdoch Children’s Research Institute, Parkville, VIC, Australia; Respiratory and Sleep Medicine, Royal Children’s Hospital, Parkville, VIC, Australia; Department of Paediatrics, University of Melbourne, Parkville, VIC, Australia; School of Mathematics and Statistics, University of Melbourne, Parkville, VIC, Australia; Centre for Cancer Research, University of Melbourne, Parkville, VIC, Australia; The Walter and Eliza Hall Institute of Medical Research, 1G Royal Parade, Parkville, VIC 3052, Australia; The Department of Medical Biology, The University of Melbourne, Parkville, VIC 3010, Australia

**Keywords:** Single-cell, Benchmarking, Demultiplexing

## Abstract

Sample multiplexing is often used to reduce cost and limit batch effects in single-cell RNA sequencing (scRNA-seq) experiments. A commonly used multiplexing technique involves tagging cells prior to pooling with a hashtag oligo (HTO) that can be sequenced along with the cells’ RNA to determine their sample of origin. Several tools have been developed to demultiplex HTO sequencing data and assign cells to samples. In this study, we critically assess the performance of seven HTO demultiplexing tools: *hashedDrops, HTODemux, GMM-Demux, demuxmix, deMULTIplex, BFF* and *HashSolo*. The comparison uses data sets where each sample has also been demultiplexed using genetic variants from the RNA, enabling comparison of HTO demultiplexing techniques against complementary data from the genetic “ground truth”. We find that all methods perform similarly where HTO labelling is of high quality, but methods that assume a bimodal counts distribution perform poorly on lower quality data. We also suggest heuristic approaches for assessing the quality of HTO counts in a scRNA-seq experiment.

## Introduction

Improvements in droplet-based single-cell RNA sequencing (scRNA-seq) technologies have prompted growing interest in exploring variation in gene expression at cellular resolution. While costs continue to decrease, it remains expensive to separately capture and sequence individual samples. Batch effects also confound meaningful differences in gene expression between samples, and robust detection of multiplets (droplets containing two or more cells) solely from the transcriptome remains an issue [17]. One solution to address these problems is to design multiplexed experiments, where samples are pooled prior to droplet capture and sequencing. The cost per sample is reduced by a factor of the number of samples sequenced, while major sample preparation batch effects within the pool are eliminated. Importantly, droplets containing cells from two or more samples can be identified. In addition, the number of cross-sample doublets can be used to estimate the expected number of within-sample doublets and thereby inform the application of other doublet detection algorithms such as *scds* [2] and *scDblFinder* [7].

Despite these advantages, it is important to carefully consider the most appropriate multiplexing protocol for the sample type(s) [5], and if additional information is required to associate the cells with their sample of origin. For genetically distinct samples, demultiplexing can be performed based on genetic variants identified from the transcriptome using a variety of tools such as *vireo* and *demuxlet* [10, 11]. However, genetic demultiplexing is not possible where samples from the same individual are sequenced together (e.g. before and after treatment or different tissues from the same individual), or in model organisms, where there is typically little genetic variation between individuals. Additionally, although genetic demultiplexing is able to distinguish cells from genetically distinct individuals, it cannot provide absolute identification of the individual sample within the pool without further information about the samples, such as SNP genotyping.

Cell hashing is an alternative multiplexing technique. Prior to pooling, a barcoded label called a hashtag oligo (HTO) is added, one to each sample. The HTOs attach to either antibodies or lipids on the surface of the cells and the HTOs are captured and sequenced in parallel to the RNA. The antibodies bind to ubiquitous cell surface proteins [19] whilst the lipids incorporate into the plasma cell membrane [15].

Sequencing of the HTOs produces a HTO counts matrix, an *N*_HTOs_ × *N*_droplets_ matrix consisting of the read counts for each HTO in each droplet. In an ideal scenario, each droplet contains only one cell and each cell contains only counts for the HTO corresponding to its sample of origin. In this ideal case, the demultiplexing algorithm involves simply identifying the non-empty entries in each column of the HTO counts matrix. In practice, the data is noisy; droplets may contain multiple cells, HTO conjugated antibodies/lipid molecules may not bind well to the cells or may dissociate and bind to cells from another sample in the pooling stage, or unbound HTOs may be present in droplets [19, 15]. Therefore, some sophistication is required for demultiplexing algorithms to distinguish the counts from the “true” HTO against a background of “false” counts.

In this study, we present a comparison of seven HTO demultiplexing methods: *hashedDrops, HTODemux, deMULTIplex, GMM-Demux, demuxmix, BFF* and *HashSolo*. We discuss the details of each in the Methods section. In all cases, the fundamental goal of each method is the same: to examine the counts of each HTO in a droplet and determine the sample of origin of each cell. Conceptually, this is achieved by separating the signal from the oligo bound to the cell in the sample preparation stage (‘positive’ HTOs) from ambient counts that arise due to contamination (‘negative’ HTOs). Droplets with no positive HTOs are classified as ‘negative’ or ‘unknown’. Droplets with more than one positive HTO are classified as doublets/multiplets. Those with only one positive HTO are classified as singlets. Here we use three data sets to assess the performance of each demultiplexing method by comparing the assignments from HTO demultiplexing to assignments from genetic demultiplexing on the same data. Recent work has shown that performance of HTOs varies between technologies and tissue types [16, 5], and the data sets herein use both antibody-derived and lipid-based HTOs and incorporate liquid and solid tissue types. Firstly, we suggest some visualisations for assessing the quality of HTO tagging. Next, we compare each method’s performance on data whose labelling quality ranges from good to poor. We find that all methods perform similarly when the labelling is of high quality. However, with lower quality labelling, methods that make simplistic, explicit assumptions about the data perform worse than those that take a more flexible approach.

## Results

### Evaluation data sets

We perform our comparison of hashtag demultiplexing methods on six tagging experiments across three data sets, each using different tagging technologies. The first data set, the BAL data set, contains 24 genetically distinct samples of bronchoalveolar lavage fluid tagged with Totalseq-A antibody-derived tags (ADTs) [19]. These samples were processed in three batches of eight pooled samples, each with two captures per batch. Batch 1 contains 24,091 droplets, batch 2 contains 48,841 droplets and batch 3, 62,306 droplets.

The second data set, the ovarian tumour (OT) data set, contains eight genetically distinct samples of high-grade serous ovarian tumours, tagged with Totalseq-B ADTs. These samples were processed in two batches of four samples each; batch 1 contains 12,510 droplets and batch 2 contains 9,547 droplets [8].

The third data set, the cell line (CL) data set, contains samples from three human lung cancer cell lines, which are tagged with MULTI-seq lipid-modified oligos (LMOs) [15]. The cell line data set contains 45,977 droplets. For each data set *vireo* [10] is used to assign cells to individuals with default settings (see Methods) and these are used as the “ground truth” to assess the accuracy of the HTO demultiplexing methods. Tests of *vireo* on simulated data sets show close to 100% accuracy on singlets and > 90% accuracy on between-sample doublets [17]. While such performance may be optimistic for real world data sets, *vireo* returns similar assignment scores for all three batches in the BAL data set (Supplementary Figure S2). This consistent performance is in contrast to the HTO demultiplexing methods.

Throughout this paper, we make a distinction between the labels assigned to cells using *vireo* and the labels of the corresponding HTOs. The *vireo* labels are denoted by the data set name followed by an alphabetical suffix e.g. BAL A, CL B, OT C, while the corresponding HTO labels contain a numeric suffix e.g. BAL 1, CL 2, OT 3.

### QC visualisation

To assess the quality of the HTO labelling and sequencing, we suggest using some common qualitative visualisations that can guide overall expectations of the performance of HTO deconvolution. Figure 1 shows the probability density function (PDF), approximated using kernel density estimation, of the logarithm of counts per cell of each HTO across the three batches in the BAL data set (Figure 1a-c). The tSNE dimensional reductions of the PCA of log-normalized HTO counts in each batch are also shown (Figure 1d-f). Each HTO in batch 1 of the BAL data set follows a bimodal distribution (Figure 1a), with a lower peak corresponding to the background counts in the majority of droplets and a higher peak corresponding to the cells from the tagged sample. In batches 2 (Figure 1b) and 3 (Figure 1c), some HTOs (e.g. BAL J and BAL N in batch 2, BAL U in batch 3) appear unimodal, indicating lower quality labelling. In the right column, the tSNE of batch 1 (Figure 1d) has 8 distinct clusters, corresponding to the 8 individual samples, with a constellation of smaller, interspersed clusters which correspond to doublets and unassigned droplets based on the genetic assignments. Batches 2 (Figure 1e) and 3 (Figure 1f) also show 8 clusters, however, the boundaries of these clusters are closer than in batch 1, and overlap for some samples in batch 3. While not quantitative, the tSNE plots in Figure 1 indicate that the cells in batch 1 are well-labelled, while those in batches 2 and 3 are labelled more poorly, highlighting that demultiplexing these batches is likely to be more challenging and demultiplexing less accurate (as shown in the following section). In addition, specific samples within a batch are labelled more poorly than others as indicated by the density plots of the individual HTOs and the overlapping tSNE clusters. Overall, the density and tSNE plots of the HTO counts can be used to quickly evaluate the quality of the HTO labelling. High quality data is indicated by bimodal density plots and tSNE plots with distinct, major clusters corresponding to the number of samples.

**Fig. 1:**
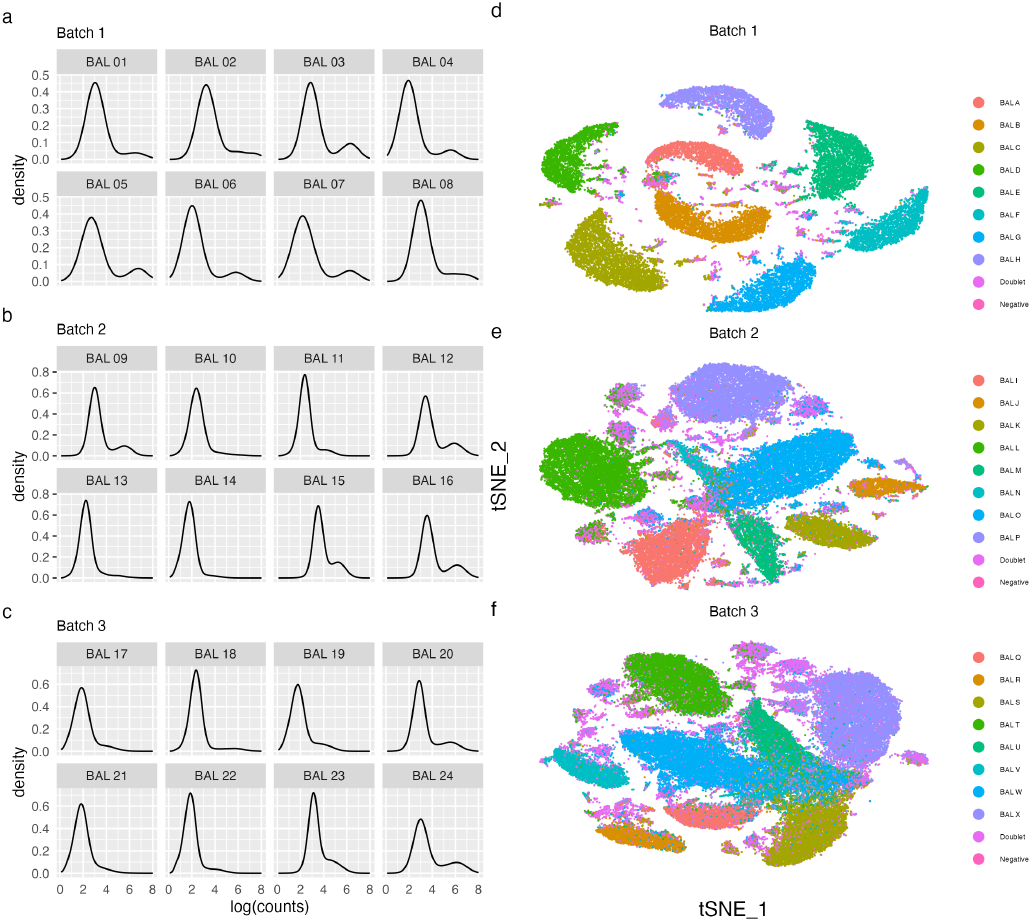
Quality assessment visualisations of the BAL data set. Left column (a-c): Probability density function of the logarithm of HTO counts for each hashtag, right column (d-f): tSNE dimensional reduction of HTO counts, coloured by genetic donor in batches 1 (d), 2 (e) and 3 (f).

### Quantitative comparisons of demultiplexing methods

Each of the three batches in our BAL data set contain cells from eight samples, from genetically distinct donors. Each demultiplexing method (including the genetic demultiplexing) can return one of 10 assignments for a cell: singlet, corresponding to one of the eight unique samples; doublet; or negative. We compare seven HTO demultiplexing methods: *BFF* (Boggy et al. 2022); *deMULTIplex* [15]; *demuxmix* [20]; *GMM-Demux* [22]; *hashedDrops* [13], *HTODemux* [19] and *HashSolo* [3]. *BFF* has two modes, *BFF*_raw_ and *BFF*_cluster_, and we present the output of both. All of the methods we consider have some adjustable parameters that affect output, however, in our exploration changing the default options does not significantly change the assignments. We discuss the details of each method and their parameters further in the Methods section. The exception is *hashedDrops*, which uses a simple counts threshold to distinguish negatives and singlets. We find that in many cases the default value of this threshold is too high, and performance (defined here as the F-score, see Methods) is improved by lowering its value. To illustrate this we present the *hashedDrops* classifications with both the default value (confident.min = 2) and the value we find maximises the F-score (confident.min = 0.5). As each batch was processed across two captures, we run the demultiplexing methods on HTO data from each individual capture. However, for simplicity, the results are presented per batch as we do not observe significant variation between captures within a batch.

In Figure 2, we show the fraction of assignments in each broad category: singlet; doublet; or negative, from *vireo* and each hashtag demultiplexing method for the three batches in the BAL data set. Two clear trends are apparent in Figure 2. First, *vireo* is able to assign more droplets as singlets than any of the hashtag demultiplexing methods. Second, the hierarchy of HTO tagging quality between the batches suggested by Figure 1 is confirmed in Figure 2. The fraction of negative droplets increases from batch 1 to batch 3 for most methods. The exception to both is *BFF*_cluster_, which assigns slightly more singlets than *vireo* in batches 1 and 2, and assigns fewer negative droplets in batches 2 and 3 than in batch 1. *HashSolo* assigns fewer negative cells than any other method, excluding *vireo* and *BFF*_cluster_, and assigns a similar fraction of doublets in each batch, with the doublet fraction greater than *vireo* in all batches.

**Fig. 2:**
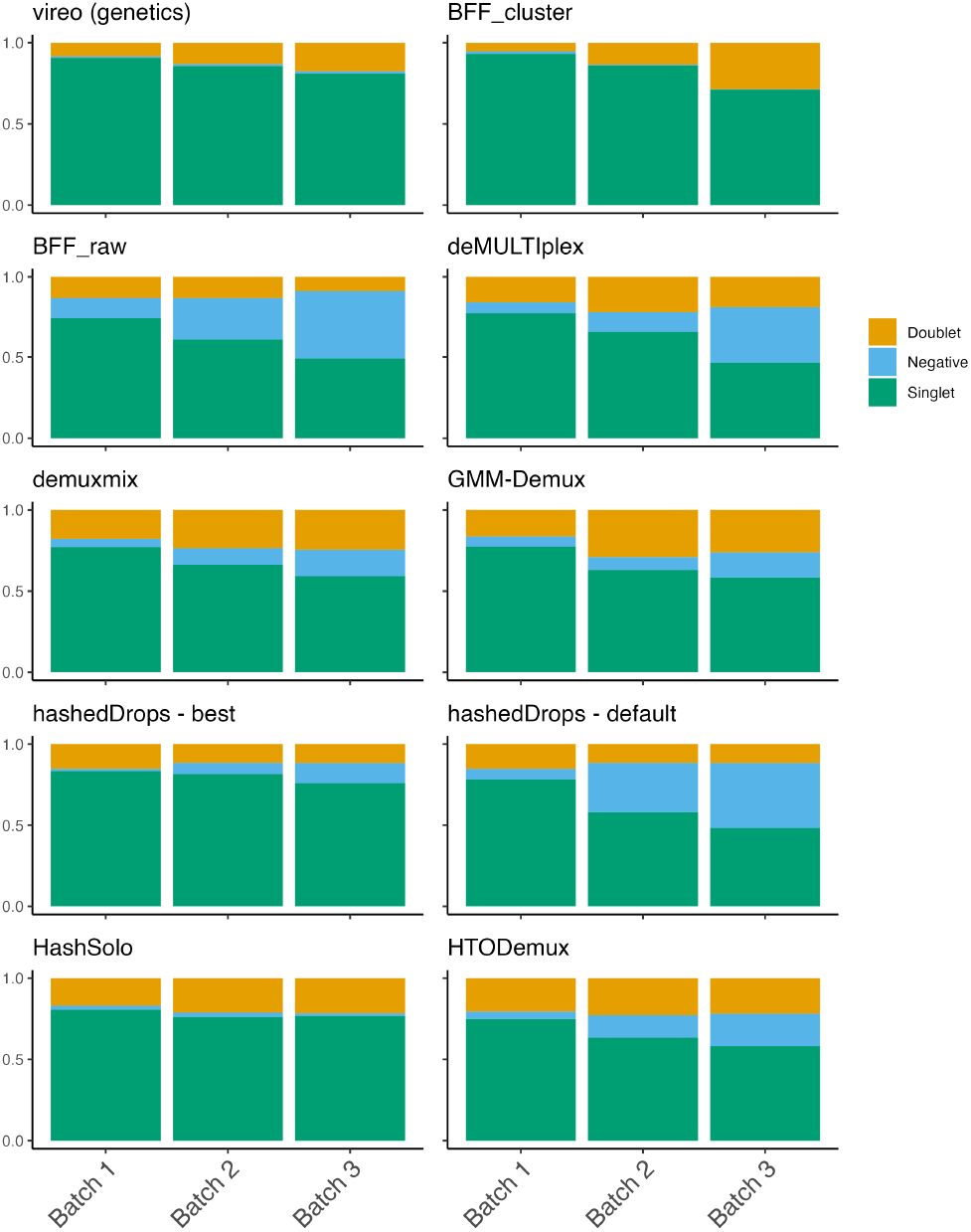
The proportion of cell assignments to singlets, doublets or negative droplets for each demultiplexing method of the BAL data set. Each panel is a method and each bar is a batch.

We next compare the specific individuals allocated by the singlet assignments of each HTO demultiplexing method to the “ground truth” of genetic assignments from *vireo*. To quantitatively assess their performance, we calculate the F-score (see Methods), a statistic which is the harmonic mean of precision and recall. The F-score ranges between 0 and 1, with a higher value indicating better performance. Figure 3 shows a heatmap of the F-score of each method, for each possible singlet assignment, split by batch.

**Fig. 3:**
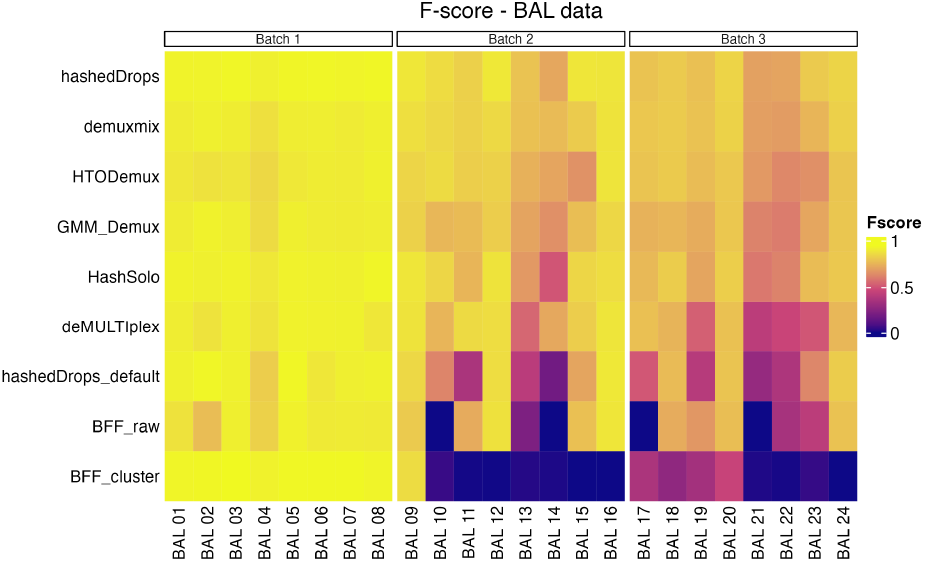
F-scores of each singlet assignment from each demultiplexing method for the BAL data set. Batch 1 (left panel), batch 2 (middle panel), batch 3 (right panel). Methods are ordered from highest to lowest average F-score across all three batches.

Table 1 shows the mean F-score of each sample for each method in all data sets. The BAL data are shown in the first three columns. Numerical values of the F-scores for all samples and methods are included in the supplementary materials. Figure 3 and Table 1 show that all methods perform well for batch 1. The overall performance of all methods drops for batches 2 and 3 and some methods begin to show significant performance differences between batches. Notably, *BFF*_cluster_, which has the highest mean F-score in batch 1, has *F* < 0.1 for all HTOs except BAL 09 in batch 2 and all HTOs except BAL 17, 18, 19, and 20 in batch 3. *BFF*_raw_ is unable to classify any cells to BAL 10 or BAL 14 in batch 2 or BAL 17, BAL 21 and HTO-14 in batch 3. *hashedDrops* has higher scores in all batches with optimised parameters than with the default settings, and has the highest mean F-score of all methods in batches 2 and 3. *Demuxmix, GMM-Demux, HashSolo* and *HTODemux* show consistent performance across all three batches.

**Table 1.** Singlet fraction and mean F-score of each demultiplexing method for all batches in all data sets. The best-performing method for each batch is indicated in bold.

| Metric | BAL Batch 1 | BAL Batch 2 | BAL Batch 3 | OT Batch 1 | OT Batch 2 | CL |
| --- | --- | --- | --- | --- | --- | --- |
| Genetic singlet fraction | 21851 / 24091 | 41816 / 48841 | 50496 / 62306 | 11296 / 12510 | 8683 / 9547 | 40118 / 45977 |
| $F_{\text{mean}}$ (*hashedDrops*) | 0.930 | **0.847** | **0.785** | **0.679** | **0.813** | 0.864 |
| $F_{\text{mean}}$ (*demuxmix*) | 0.907 | 0.833 | 0.775 | 0.511 | 0.506 | 0.743 |
| $F_{\text{mean}}$ (*HTODemux*) | 0.892 | 0.794 | 0.742 | 0.620 | 0.654 | 0.737 |
| $F_{\text{mean}}$ (*GMM-Demux*) | 0.905 | 0.770 | 0.720 | 0.645 | 0.632 | 0.823 |
| $F_{\text{mean}}$ (*HashSolo*) | 0.920 | 0.783 | 0.737 | 0.606 | 0.688 | 0.806 |
| $F_{\text{mean}}$ (*deMULTiplex*) | 0.907 | 0.791 | 0.624 | 0.218 | 0.684 | 0.828 |
| $F_{\text{mean}}$ (*hashedDrops* - default) | 0.907 | 0.613 | 0.573 | 0.406 | 0.46 | 0.663 |
| $F_{\text{mean}}$ (*BFF<sub>raw</sub>*) | 0.879 | 0.539 | 0.466 | 0 | 0.623 | 0.791 |
| $F_{\text{mean}}$ (*BFF<sub>cluster</sub>*) | **0.935** | 0.119 | 0.188 | 0 | 0.408 | **0.874** |

Next, we investigate doublets in more detail. As shown in Figure 2, almost all methods assign more doublets than *vireo*. Assigning true doublets as singlets is potentially a more significant source of error in downstream analysis, such as cell type identification, than misclassifying a true singlet as a doublet, negative, or incorrect singlet [21]. Therefore, we exclude the doublet and negative classifications from our F-score analysis above. Instead, we perform a separate, complementary analysis of how each HTO demultiplexing method classifies the genetic doublets. For each batch in the BAL data set, we take the droplets assigned as doublets by *vireo*, and look at which broad category (i.e. doublet, singlet or negative) they are assigned by each of the HTO demultiplexing methods. For this analysis, the best performing method is the one which minimizes the number of “true” doublets assigned as singlets. Since both negative droplets and doublets are typically excluded from downstream analysis, misclassification of genetic doublets as negatives is relatively unimportant.

Figure 4 shows the fraction of *vireo* doublets assigned by each method as either doublet, singlet or negative. Figure 4 illustrates several points not apparent in Figure 3 and Table 1. Firstly, *BFF*_cluster_, which has the highest F-score for batch 1, has the worst performance on the doublets, assigning more than half of the genetic doublets in that batch as singlets. Secondly, while adjusting the parameters of *hashedDrops* from their default values improves the F-score, the number of incorrectly assigned genetic doublets approximately doubles in all batches. Thirdly, the other best-performing methods based on F-score: *demuxmix, HTODemux* and *GMM-demux*, perform well on the doublet analysis as well, assigning < 20% of genetic doublets as singlets in all batches, though *HashSolo* performs slightly worse, with ≈ 30% of genetic doublets identified as singlets.

**Fig. 4:**
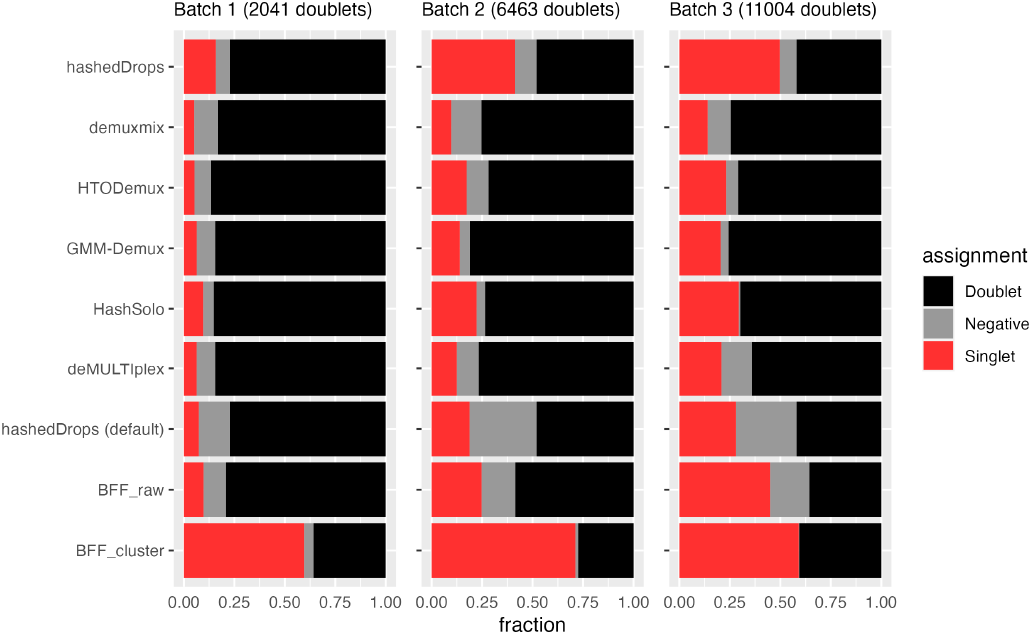
Fraction of genetic doublets assigned by each HTO demultiplexing method to doublets (black), negatives (grey) or singlets (red) in the BAL data set. Methods are in the same order as Figure 3.

### Ovarian tumour data

The second data set we analyse consists of eight samples from high-grade serous ovarian carcinoma patients from [8]. Unlike the BAL data samples, which are liquid, these tumour samples require dissociation prior to hashtagging and single-cell sequencing. The samples were tagged with Totalseq-B ADTs and processed in two batches of four samples each, with 12,510 droplets in batch 1 and 9,547 droplets in batch 2.

Genetic demultiplexing with *vireo* positively assigns > 90% of droplets as singlets for both batches [8]. Initial QC of the HTO counts (Supplementary Figure S6) suggests poor performance of the hashtagging in this experiment, which is borne out by the low F-scores of all demultiplexing methods on this data set.

Figure 5 shows a heatmap of the F-scores for each method on each sample and the assignment of genetic doublets as either doublets, singlets or negative by each method. The mean F-scores for each method in each batch are shown in the fourth and fifth columns of Table 1. In general, the performance of all demultiplexing methods is worse for both batches in the ovarian tumour data set than the BAL data. This is possibly due to the additional dissociation step in the sample processing [5]. As for the BAL data set, using *hashedDrops* is a trade-off between F-score and doublet classification accuracy. *HashSolo, GMM-Demux* and *HTODemux* all perform relatively well, while *demuxmix*, the method with best overall performance on the BAL data set, falls behind on F-score but makes fewer errors on doublets. How these two factors should be weighed against one another cannot be answered objectively for all cases, especially as the overall fraction of doublets depends on the overall number of cells per batch [17]. For example, in batch 3 of the BAL data set, using *hashedDrops* instead of *demuxmix* improves the mean F-score by only 0.01, while an additional ≈ 3000 doublets are incorrectly labelled as singlets. In batch 1 of the ovarian tumour data set, however, using *hashedDrops* instead of *demuxmix* improves the mean F-score by 0.307, while misclassifying ≈ 500 doublets as singlets. In the former case *demuxmix* is clearly superior, while in the latter case it might make sense to prefer *hashedDrops. BFF*_raw_ and *BFF*_cluster_ again perform poorly, especially on batch 1, where both fail to classify any cells.

**Fig. 5:**
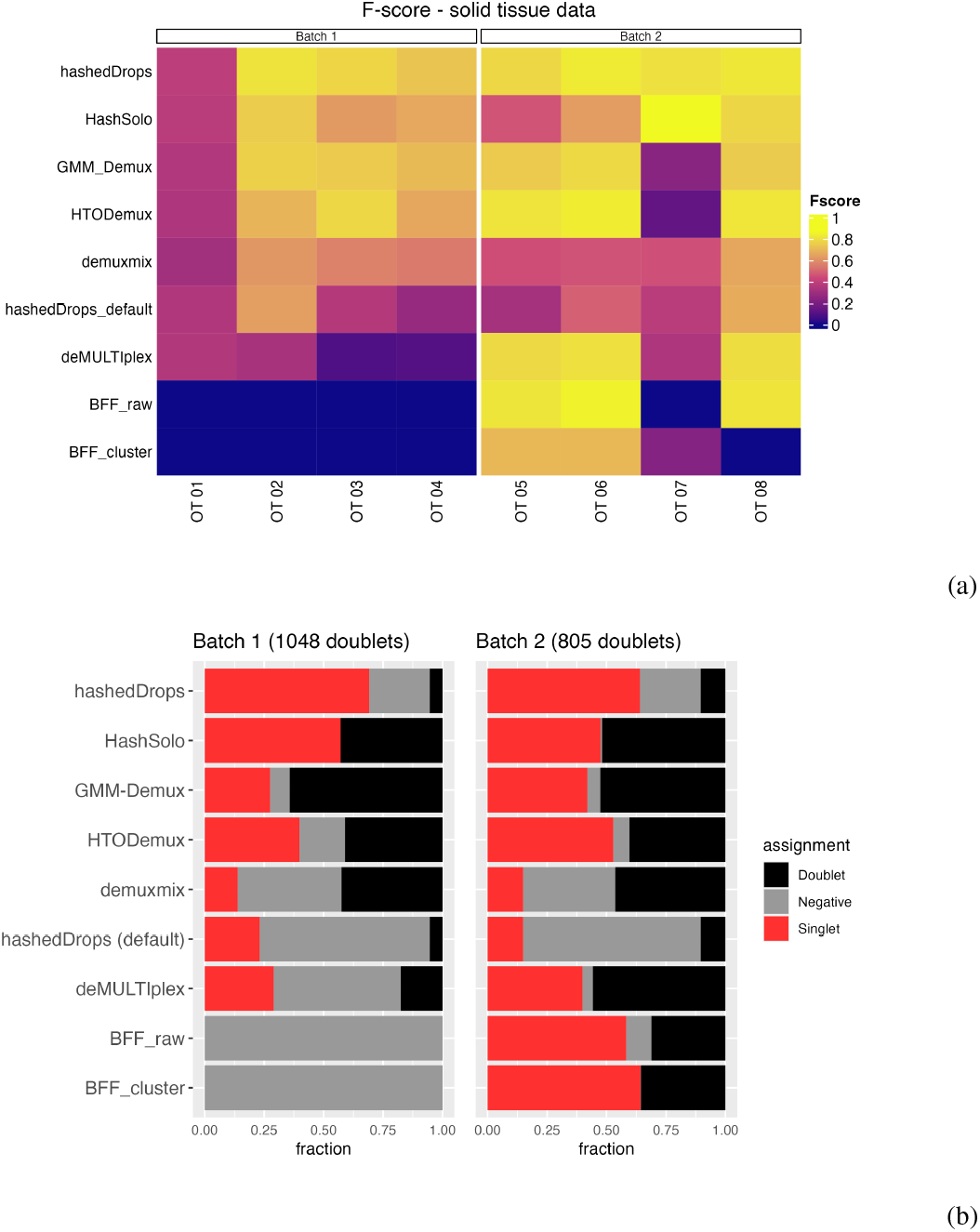
(a) Heatmap of F-scores for each demultiplexing method on each sample from the ovarian tumour data set. (b) Fraction of the genetic doublets in each batch assigned to different categories by each method. Methods are ordered from highest to lowest average F-score across the two batches.

### Cell line data

We perform the same analysis on a third data set, the cell line data set, consisting of three samples from genetically distinct human lung cancer cell lines. Here, H3122, H358 and H1792 cells were tagged with different MULTI-seq LMOs [15], a different tagging technology to the ADTs used on the BAL and ovarian tumour samples. These samples were pooled together and processed in one batch across three captures, with 45,977 total droplets.

Since both the ADT and LMO technologies produce an *N*_HTOs_ × *N*_droplets_ counts matrix with similar distributions (see supplementary Figure S7), we expect the demultiplexing methods to perform similarly on the LMO and ADT data. Figure 6 shows the F-score for each method on each of the three samples in this data set (Figure 6a), as well as the categorical assignments of the 4945 genetic doublets (Figure 6b). Figure 6 and Figure S7 show that the cell line data is somewhere between the quality of batches 1 and 2 of the BAL data, and the performance of each of the methods is similar. Based on F-score alone (Figure 5a), *BFF*_cluster_ performs best, however, looking at Figure 5b we see that more than 75% of genetic doublets are assigned as singlets. Based on the two metrics, we find that *deMULTIplex, GMM-Demux* and *demuxmix* perform well, *hashedDrops* with default parameters and *HTODemux* perform relatively poorly, and *hashedDrops* with lowered thresholds and *HashSolo* perform well based on the F-score, but misidentify nearly 60% of genetic doublets as singlets – more than twice as many as the best-performing methods.

**Fig. 6:**
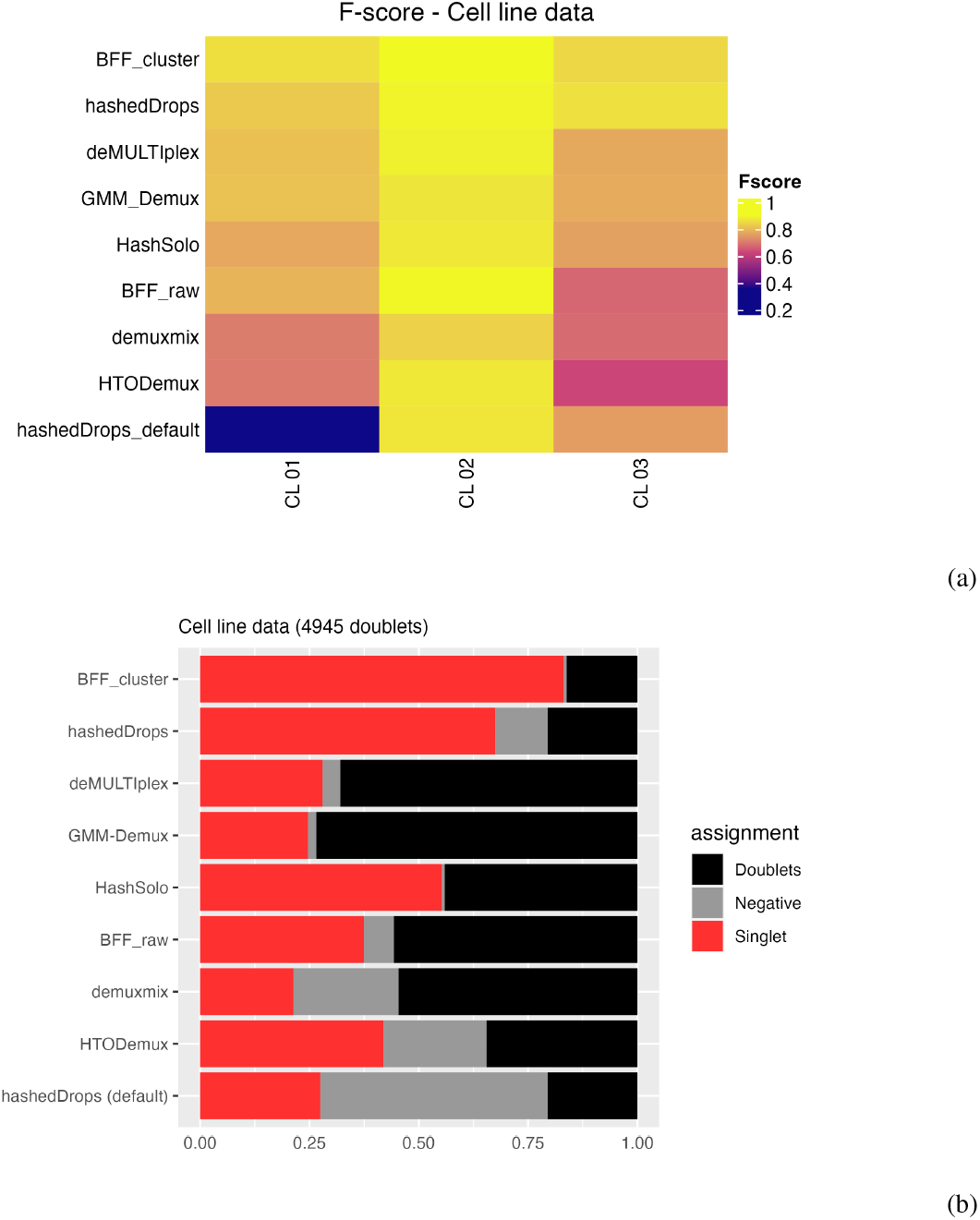
(a) Heatmap of F-scores for each demultiplexing method on each sample from the cell line data set. (b) Fraction of the genetic doublets assigned to different categories by each method. Methods are ordered from highest to lowest average F-score.

## Discussion

As sample multiplexing becomes more common in scRNA-seq experiments, reliable demultiplexing of cells becomes paramount. We benchmark seven methods for cell demultiplexing based on hashtag oligo data. Of the methods we consider, *demuxmix* shows the best overall performance across the three data sets included in this study, using our two criteria of accurately classifying singlets and rarely misclassifying genetic doublets as singlets. However, the difference between *demuxmix, GMM-Demux* and *HTODemux* is small, and all should perform relatively well on most data sets. Furthermore, for data sets with lower-quality hashing, we suggest that it may be prudent to trial several of these methods to maxmise the number of positively-identified singlets. Looking only at the F-score, *hashedDrops* is the best overall performer when the threshold for confident detection is lowered. However, this comes at the cost of misclassifying doublets as singlets. *deMULTIplex* and the *BFF* methods perform especially poorly on lower-quality hashing data, as they rely on measurement of two peaks in a density estimate of the transformed counts, which may not exist when a large number of background counts are present.

Our results are broadly consistent for hashtagging using ADTs and LMOs, as well as liquid and solid tissues, indicating that the performance of the demultiplexing methods is agnostic to the choice of tagging protocol.

Although most of the tools are straightforward to run, and interact well with popular single-cell analysis packages, there are some important usability differences. *Demuxmix* is part of the *Bioconductor* ecosystem and can easily be run in R. As it only requires a HTO counts matrix to return assignments it can be incorporated as part of a *Bioconductor* or *Seurat*-based single-cell analysis pipeline. *HTODemux* is part of the *Seurat* package and requires a *Seurat* object as input, therefore runs most easily alongside other *Seurat* tools for single-cell analysis. *HashSolo* is part of the *scanpy* ecosystem and can be run easily as part of a *scanpy* analysis pipeline. However, although *HashSolo* performs well on high quality data, its tendency to misidentify genetic doublets as singlets means care should be taken on super-loaded data. *GMM-Demux* is a command-line tool, which may provide a barrier to entry for some users, although wrappers such as *cellhashR* [4] can be used to run it from R.

We demonstrate two simple visualisation methods to assess the quality of hashtag data, and confirm that if the probability density of counts follows a bimodal distribution, and the counts separate into well-defined clusters on a dimensional reduction plot, then all demultiplexing methods perform well. However, if these conditions are not met, demultiplexing algorithms that explicitly assume bimodal distributions (such as *deMULTIplex* and *BFF*) fail to correctly assign some droplets to their samples of origin. Threshold-based methods, such as *hashedDrops*, can perform well but make a trade-off between greater recovery of singlets and false positives. More sophisticated methods, such as the clustering-based *HTODemux* and *demuxmix*, and the Bayesian estimation-based methods *GMM-Demux* and *Hashsolo* perform best and most consistently on both high and low-quality hashtag data.

Low-quality hashtag data does not imply low-quality RNA expression data; importantly, the two are largely uncorrelated (see Figure S1 in Supplementary Materials). We show that the difference between demultiplexing methods becomes more pronounced as the quality of the hashtag data reduces. Therefore, maximising performance of demultiplexing methods on lower-quality hashtag data is particularly important to prevent otherwise good quality cells being excluded in a single-cell analysis.

For samples with similar or identical genetic backgrounds, labelling of cells before pooling will continue to be an important strategy and this manuscript provides a benchmark for applying these methods. While genetic demultiplexing has been shown to be accurate for deconvolving samples from out-bred populations [17], there is potential for information from the genetics and hashtags to be used together for more accurate deconvolution [6, 12] and we expect to see continued developments in this area.

## Methods

### Single-cell data generation

The BAL data set is derived from CITE-seq experiments of 24 samples of paediatric BAL. Samples were collected, cryopreserved, and thawed as previously described [18]. Live, single cells were sorted using a BD FACS Aria fusion and resuspended in 25μL of cell staining buffer (BioLegend). Human TruStain FcX FC blocking reagent (BioLegend) was added according to manufacturers’ instructions for 10min on ice. Each tube was made up to 100μL with cell staining buffer and TotalSeq Hashtag reagents (BioLegend) were added to each sample for 20min on ice. Cells were washed with 3mL cell staining buffer and centrifuged at 400xg for 5min at 4°C. Supernatant was discarded and each sample resuspended at 62,500 cells/100μL following which 100μL of each sample were pooled into one tube. Pooled cells were centrifuged at 400xg for 5min at 4°C, supernatant discarded, and resuspended in 25μL cell staining buffer and 25ul of TotalSeqA Human Universal Cocktail v1.0 (BioLegend) for 30min on ice. This cocktail contains 154 immune related surface proteins. Cells were washed in 3mL cell staining buffer and centrifuged at 400xg for 5min at 4°C. Following two more washes, cells were resuspended in PBS + 0.04% BSA for Chromium captures. Single-cell captures, library preparation, and sequencing was performed as we have described previously [14].

Data for the ovarian tumour data set was taken from [8], and was provided as a matrix of HTO counts along with the genetic assignments from vireo.

For the cell line data set, three human lung cancer cell lines: H1792, H3122 and H358 were labelled with a different 3’ lipid modified oligo (LMO) as in [15]. Cell lines were pooled in a 1:1:1 ratio and the pool was used for three separate captures with the 10x Chromium system using the 10X Genomics NextGEM 3’ Single-cell Gene Expression Solution (10x Genomics). Post single-cell capture, scRNA libraries were generated according to the manufacturer’s recommendations and LMO library preparation was performed as described previously [15]. LMO count matrices were generated from fastq files using *CITE-seq-count* v 1.4.3.

### Genetic demultiplexing

For both data sets, genetic donors were assigned to the samples by first performing SNP genotyping using *cellSNP-lite* (v1.2.0 for the BAL data; v1.2.1 for the cell line data) [9]. We used a list of common variants from the 1000 genome project [1] and filtered SNPs with < 20 UMIs or < 10% minor alleles, as recommended in the *cellSNP-lite* manual. We then used *vireoSNP* 0.5.6 [10] for demultiplexing using the output of *cellSNP-lite* as the cell data and no additional donor information. More detail is provided in [14].

### Calculating the F-score

For each possible HTO assignment we calculate the true positive rate TP, which is the fraction of cells with that HTO assignment that have the corresponding *vireo* assignment; the false positive rate FP, which is the fraction of cells with that HTO assignment and a different genetic assignment; and the false negative rate FN, which is the fraction of cells with the corresponding genetic assignment but a different HTO assignment. Our key metric is the F-score, which is defined as:

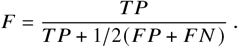

F is the harmonic mean of the precision and recall, and can vary between 0 and 1, with a higher F-score implying better performance.

### Overview of demultiplexing methods in this comparison

#### hashedDrops

*hashedDrops*, part of the *DropletUtils* package [13], is a simple threshold-based classifier. First, the HTO counts matrix is corrected for the ambient counts of each HTO in the data (either before or after filtering out empty droplets). It then ranks the HTO counts in each droplet. Assignments are determined solely by the log-fold change (LFC) between the highest and second highest counts in a droplet, relative to the median counts for that HTO. Firstly, doublets are called where the LFC of the second highest HTO is greater than a user-defined number of median absolute deviations (MAD) above the median and also greater than another user-defined threshold. If a droplet is not assigned as a doublet, singlet assignments are determined by checking that the LFC of the HTO with the highest count in each droplet is greater than a user-defined threshold and is also not less than a user-defined number of MADs below the median. While less sophisticated than other demultiplexing methods, hashedDrops has the advantage of making very few assumptions about the data, and is easily configurable by the user. However, as the results are very sensitive to the choice of the hard thresholds their values should be carefully considered. We explore the effect of varying the singlet threshold parameter in Figure S3 in the supplementary materials.

#### HTODemux

*HTODemux* [19], included in the *Seurat* package, uses a clustering-based approach. The HTO counts are normalized using the centred log ratio (CLR) transformation. Then an unsupervised *k*-medoids clustering is performed, with *k* = *N*_HTOs_ + 1. For each HTO, cells are identified as positive or negative in a two step procedure. Firstly, the cluster with the lowest expression count for each HTO is defined as the “negative” cluster, and a negative binomial distribution is fitted to the counts in that cluster. For the droplets outside that cluster, droplets with HTO counts above a user-defined quantile (0.99 by default) are assigned as positive for the HTO. After performing this procedure on all HTOs, droplets that have been assigned positive for more than one HTO are classified as multiplets, droplets with no positive assignments are classified as negative, and the droplets assigned positive for only one HTO are classified as singlets. We explore the effect of varying the quantile threshold in Figure S4 in the supplementary materials.

#### GMM-Demux

Like *HTODemux, GMM-Demux* [22] uses the CLR-transformed HTO counts. In well-behaved data, the distribution of the CLR-transformed counts of each HTO is bimodal, with the lower peak corresponding to the ‘negative’ background and the higher peak corresponding to the true ‘positive’ counts. *GMM-Demux* fits a two-component Gaussian mixture model to the distribution of each HTO, and uses Bayesian estimation to assign each droplet to the higher-or lower-peaked distribution for each HTO. Droplets with only one positive HTO assignment are classified as singlets, droplets with no positive assignments are classified as negative, while droplets with multiple positive assignments are classified as multiplets, with the identity of the most-probable HTOs in each multiplet included in the output. Every positive assignment is given a confidence score between 0 and 1, and a user-defined confidence threshold (0.8 by default) can be adjusted to be more or less strict with the output classifications. We explore the effect of varying the confidence threshold set in Figure S5 in the supplementary materials.

#### demuxmix

*demuxmix* [20] is similar to *GMM-Demux* but uses a negative binomial mixture model on the untransformed HTO counts, rather than a mixed Gaussian on the CLR-transformed counts. For each HTO, all cells are clustered into positive and negative clusters using *k*-means clustering. Cells with very high counts are marked as outliers, and the non-outliers are fitted to a two-component negative binomial distribution using an expectation-maximisation algorithm. *demuxmix* can also leverage the RNA counts to improve performance, using the number of detected genes in the RNA library as a covariate in the mixture model. Using this additional RNA information with the BAL data set showed no significant improvement on either data set in this paper, so the results presented are based on the HTO counts only.

#### deMULTIplex

*deMULTIplex* [15] uses an iterative approach. First, a kernel density estimator is used to smooth the log-normalized HTO counts. For each HTO, an initial threshold for positive classification is defined as the highest maximum (assuming a bimodal normalized counts distribution), while the initial threshold for negative classification is the mode. Then, the algorithm sweeps through the quantile range between these two thresholds to find the value that classifies the largest proportion of the data as singlets. Each droplet is then compared against each HTO-specific threshold, being classified as negative, singlet or multiplet based on the number of HTOs for which it passes. All negatively-classified droplets are removed from the counts matrix, and the process is repeated until successive iterations identify no additional negative droplets. While the thresholds for singlets and doublets can be adjusted manually, the default option searches for the value which maximises the fraction of singlet assignments, and our results use this automatic threshold-determining mode.

#### BFF

*Bimodal Flexible Fitting* (*BFF*) [4] also assumes a bimodal counts distribution. It operates in two modes, *BFF*_raw_ and *BFF*_cluster_. The first, *BFF*_raw_ smooths the counts distribution using a kernel density estimator, much like *deMULTIplex*. The threshold between positive and negative classification in this case is the local minimum between the two peaks. The second mode, *BFF*_cluster_, is similar, but includes an additional layer of normalization, called bimodal quantile normalization, before finalizing classifications. The level of smoothing on the counts can be selected by the user, however our results are based on the default, which searches for an optimal value.

#### HashSolo

*HashSolo* [3] is a Bayesian method that models the overall counts distribution across all cells as a mixture of two log-normal distributions corresponding to signal and noise. For each cell, it looks at the two highest counts and computes the likelihood of both belonging to the noise distribution (negative), one belonging to the signal and one the noise (singlet) or both belonging to the signal (doublet). It then returns the assignment with the highest Bayesian evidence. The prior is the fraction of singlets, doublets and negative cells within the sample. Based on the *vireo* results, we use a prior of 75% singlets, 20% doublets and 5% negative. We also run *HashSolo* with a negative fraction prior between 1% and 10% and a doublet fraction prior between 10% and 30%. This has a negligible effect on the posterior assignments, with the average F-score varying by < 1% in all batches.

## Data availability

Code to reproduce the analysis in this paper can be viewed here https://oshlacklab.com/hashtag-demux-paper, and the raw counts data and *vireo assignments* for the BAL and cell line data sets can be accessed at https://github.com/Oshlack/hashtag-demux-paper. The ovarian tumour data set is from [8], and can be accessed at https://www.ncbi.nlm.nih.gov/projects/gap/cgi-bin/study.cgi?study_id=phs002262.v2.p1

## Acknowledgments

We thank WEHI Advanced Genomics Facility (Casey Anttila, Ling Ling and Daniela Zalcenstein for cell capture and library preparation, Sarah MacRaild and Stephen Wilcox for sequencing). We also thank Anna Trigos for helpful discussion. This work was supported by CZI Inflammation grant DAF2020-217531, AO NHMRC Ideas Grant GNT1187748 and Investigator Grant GNT1196256.

## References

1. 1000 Genomes Project Consortium, Adam Auton, Lisa D Brooks, Richard M Durbin, Erik P Garrison, Hyun Min Kang, Jan O Korbel, Jonathan L Marchini, Shane McCarthy, Gil A McVean, and Gonçalo R Abecasis. A global reference for human genetic variation. Nature, 526(7571):68–74, October 2015.

2. Abha S Bais and Dennis Kostka. scds: computational annotation of doublets in single-cell RNA sequencing data. Bioinformatics, 36(4):1150–1158, February 2020.

3. Nicholas J Bernstein, Nicole L Fong, Irene Lam, Margaret A Roy, David G Hendrickson, and David R Kelley. Solo: Doublet identification in Single-Cell RNA-Seq via Semi-Supervised deep learning. Cell Syst, 11(1):95–101.e5, July 2020.

4. Gregory J Boggy, G W McElfresh, Eisa Mahyari, Abigail B Ventura, Scott G Hansen, Louis J Picker, and Benjamin N Bimber. BFF and cellhashr: analysis tools for accurate demultiplexing of cell hashing data. Bioinformatics, 38(10):2791–2801, May 2022.

5. Daniel V Brown, Casey J A Anttila, Ling Ling, Patrick Grave, Tracey M Baldwin, Ryan Munnings, Anthony J Farchione, Vanessa L Bryant, Amelia Dunstone, Christine Biben, Samir Taoudi, Tom S Weber, Shalin H Naik, Anthony Hadla, Holly E Barker, Cassandra J Vandenberg, Genevieve Dall, Clare L Scott, Zachery Moore, James R Whittle, Saskia Freytag, Sarah A Best, Anthony T Papenfuss, Sam W Z Olechnowicz, Sarah E MacRaild, Stephen Wilcox, Peter F Hickey, Daniela Amann-Zalcenstein, and Rory Bowden. A risk-reward examination of sample multiplexing reagents for single cell RNA-Seq. June 2023.

6. Fabiola Curion, Xichen Wu, Lukas Heumos, Mariana Gonzales, Lennard Halle, Melissa Grant-Peters, Charlotte Rich-Griffin, Hing-Yuen Yeung, Calliope A Dendrou, Herbert B Schiller, and Fabian J Theis. hadge: a comprehensive pipeline for donor deconvolution in single cell. August 2023.

7. Pierre-Luc Germain, Aaron Lun, Carlos Garcia Meixide, Will Macnair, and Mark D Robinson. Doublet identification in single-cell sequencing data using scdblfinder. F1000Res., 10:979, September 2021.

8. Ariel A Hippen, Dalia K Omran, Lukas M Weber, Euihye Jung, Ronny Drapkin, Jennifer A Doherty, Stephanie C Hicks, and Casey S Greene. Performance of computational algorithms to deconvolve heterogeneous bulk tumor tissue depends on experimental factors. January 2023.

9. Xianjie Huang and Yuanhua Huang. Cellsnp-lite: an efficient tool for genotyping single cells. Bioinformatics, May 2021.

10. Yuanhua Huang, Davis J McCarthy, and Oliver Stegle. Vireo: Bayesian demultiplexing of pooled single-cell RNA-seq data without genotype reference. Genome Biol., 20(1):273, December 2019.

11. Hyun Min Kang, Meena Subramaniam, Sasha Targ, Michelle Nguyen, Lenka Maliskova, Elizabeth McCarthy, Eunice Wan, Simon Wong, Lauren Byrnes, Cristina M Lanata, Rachel E Gate, Sara Mostafavi, Alexander Marson, Noah Zaitlen, Lindsey A Criswell, and Chun Jimmie Ye. Multiplexed droplet single-cell RNA-sequencing using natural genetic variation. Nat. Biotechnol., 36(1):89–94, January 2018.

12. Lei Li, Jiayi Sun, Yanbin Fu, Siriruk Changrob, Joshua J C McGrath, and Patrick C Wilson. A hybrid single cell demultiplexing strategy that increases both cell recovery rate and calling accuracy. April 2023.

13. Aaron T L Lun, Samantha Riesenfeld, Tallulah Andrews, The Phuong Dao, Tomas Gomes, participants in the 1st Human Cell Atlas Jamboree, and John C Marioni. EmptyDrops: distinguishing cells from empty droplets in droplet-based single-cell RNA sequencing data. Genome Biol., 20(1):63, March 2019.

14. Jovana Maksimovic, Shivanthan Shanthikumar, George Howitt, Peter F Hickey, William Ho, Casey Anttila, Daniel V Brown, Anne Senabouth, Dominik Kaczorowski, Daniela Amann-Zalcenstein, Joseph E Powell, Sarath C Ranganathan, Alicia Oshlack, and Melanie R Neeland. Multimodal single cell analysis of the paediatric lower airway reveals novel immune cell phenotypes in early life health and disease. June 2022.

15. Christopher S McGinnis, David M Patterson, Juliane Winkler, Daniel N Conrad, Marco Y Hein, Vasudha Srivastava, Jennifer L Hu, Lyndsay M Murrow, Jonathan S Weissman, Zena Werb, Eric D Chow, and Zev J Gartner. MULTI-seq: sample multiplexing for single-cell RNA sequencing using lipid-tagged indices. Nat. Methods, 16(7):619–626, July 2019.

16. Viacheslav Mylka, Irina Matetovici, Suresh Poovathingal, Jeroen Aerts, Niels Vandamme, Ruth Seurinck, Kevin Verstaen, Gert Hulselmans, Silvie Van den Hoecke, Isabelle Scheyltjens, Kiavash Movahedi, Hans Wils, Joke Reumers, Jeroen Van Houdt, Stein Aerts, and Yvan Saeys. Comparative analysis of antibody- and lipid-based multiplexing methods for single-cell RNA-seq. Genome Biol., 23(1):55, February 2022.

17. Drew Neavin, Anne Senabouth, Jimmy Tsz Hang Lee, Aida Ripoll, sc-eQTLGen Consortium, Lude Franke, Shyam Prabhakar, Chun Jimmie Ye, Davis J McCarthy, Marta Melé, Martin Hemberg, and Joseph E Powell. Demuxafy: Improvement in droplet assignment by integrating multiple single-cell demultiplexing and doublet detection methods. March 2022.

18. Shivanthan Shanthikumar, Matthew Burton, Richard Saffery, Sarath C Ranganathan, and Melanie R Neeland. Single-Cell flow cytometry profiling of BAL in children. Am. J. Respir. Cell Mol. Biol., 63(2):152–159, August 2020.

19. Marlon Stoeckius, Shiwei Zheng, Brian Houck-Loomis, Stephanie Hao, Bertrand Z Yeung, William M Mauck, 3rd, Peter Smibert, and Rahul Satija. Cell hashing with barcoded antibodies enables multiplexing and doublet detection for single cell genomics. Genome Biol., 19(1):224, December 2018.

20. John F Tuddenham, Mariko Taga, Verena Haage, Tina Roostaei, Charles White, Annie Lee, Masashi Fujita, Anthony Khairallah, Gilad Green, Bradley Hyman, Matthew Frosch, Sarah Hopp, Thomas G Beach, John Corboy, Naomi Habib, Hans-Ulrich Klein, Rajesh Kumar Soni, Andrew F Teich, Richard A Hickman, Roy N Alcalay, Neil Shneider, Julie Schneider, Peter A Sims, David A Bennett, Marta Olah, Vilas Menon, and Philip L De Jager. A cross-disease human microglial framework identifies disease-enriched subsets and tool compounds for microglial polarization. June 2022.

21. Samuel L Wolock, Romain Lopez, and Allon M Klein. Scrublet: Computational identification of cell doublets in Single-Cell transcriptomic data. Cell Syst, 8(4):281–291.e9, April 2019.

22. Hongyi Xin, Qiuyu Lian, Yale Jiang, Jiadi Luo, Xinjun Wang, Carla Erb, Zhongli Xu, Xiaoyi Zhang, Elisa Heidrich-O’Hare, Qi Yan, Richard H Duerr, Kong Chen, and Wei Chen. GMM-Demux: sample demultiplexing, multiplet detection, experiment planning, and novel cell-type verification in single cell sequencing. Genome Biol., 21(1):188, July 2020.

