## Supplementary materials for "Benchmarking single-cell hashtag oligo demultiplexing methods"

### RNA quality metrics

In this study we use the genetic assignment of cells to samples as a "ground truth" against which we compare the performance of hashtag demultiplexing methods. This implicitly assumes that the RNA counts are independent of the corresponding hashtag counts in each droplet. Here we show some evidence supporting this assumption, and demonstrate that genetic demultiplexing with *vireo* performs more consistently between batches than any of the hashtag demultiplexing methods.

Firstly, we look at the correlation between the RNA counts and HTO counts for each batch in the BAL data set in Figure S1.

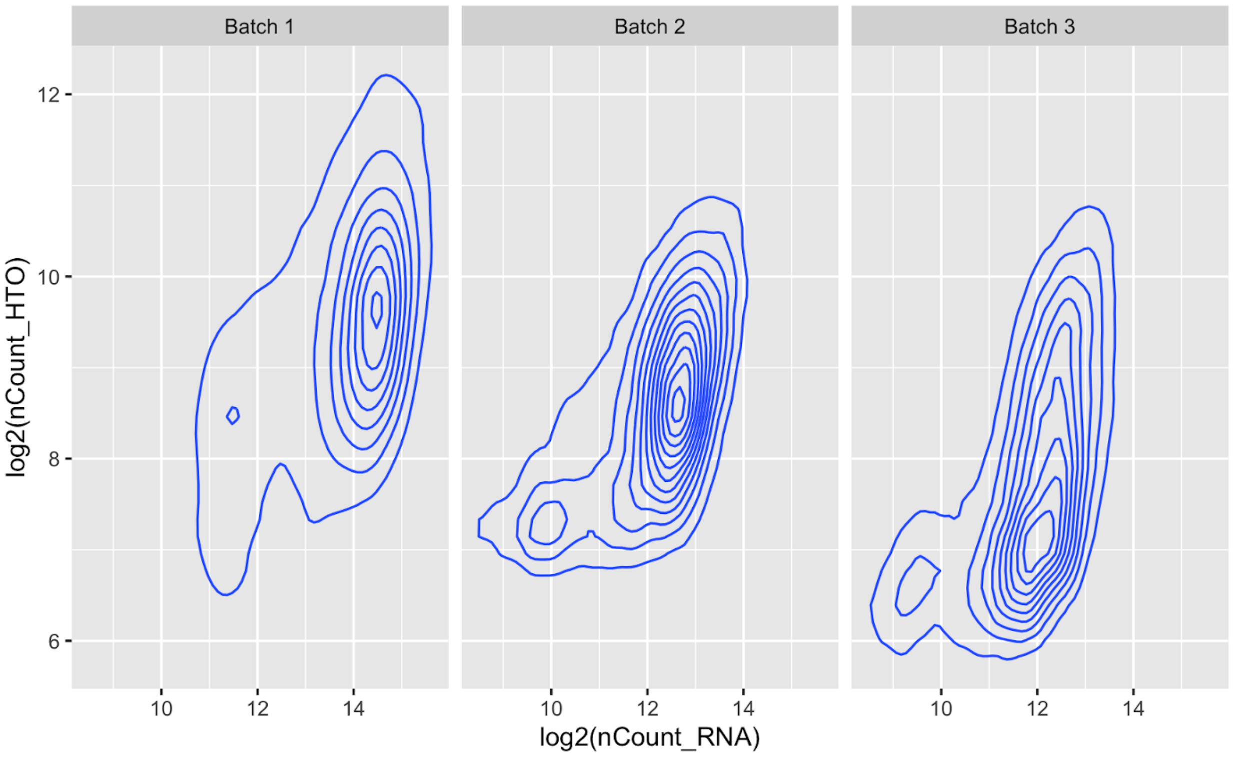


Figure S1: Contour plots of RNA counts and HTO counts from batches 1, 2 and 3 of the first data set.

Figure S1 shows that there is little correlation between the RNA and HTO counts (Pearson correlation coefficients 0.23, 0.08 and 0.27 for batches 1, 2 and 3 respectively).

Next, we look at the confidence score of singlet assignments from *vireo* for each of the three batches in the first data set in Figure S2.

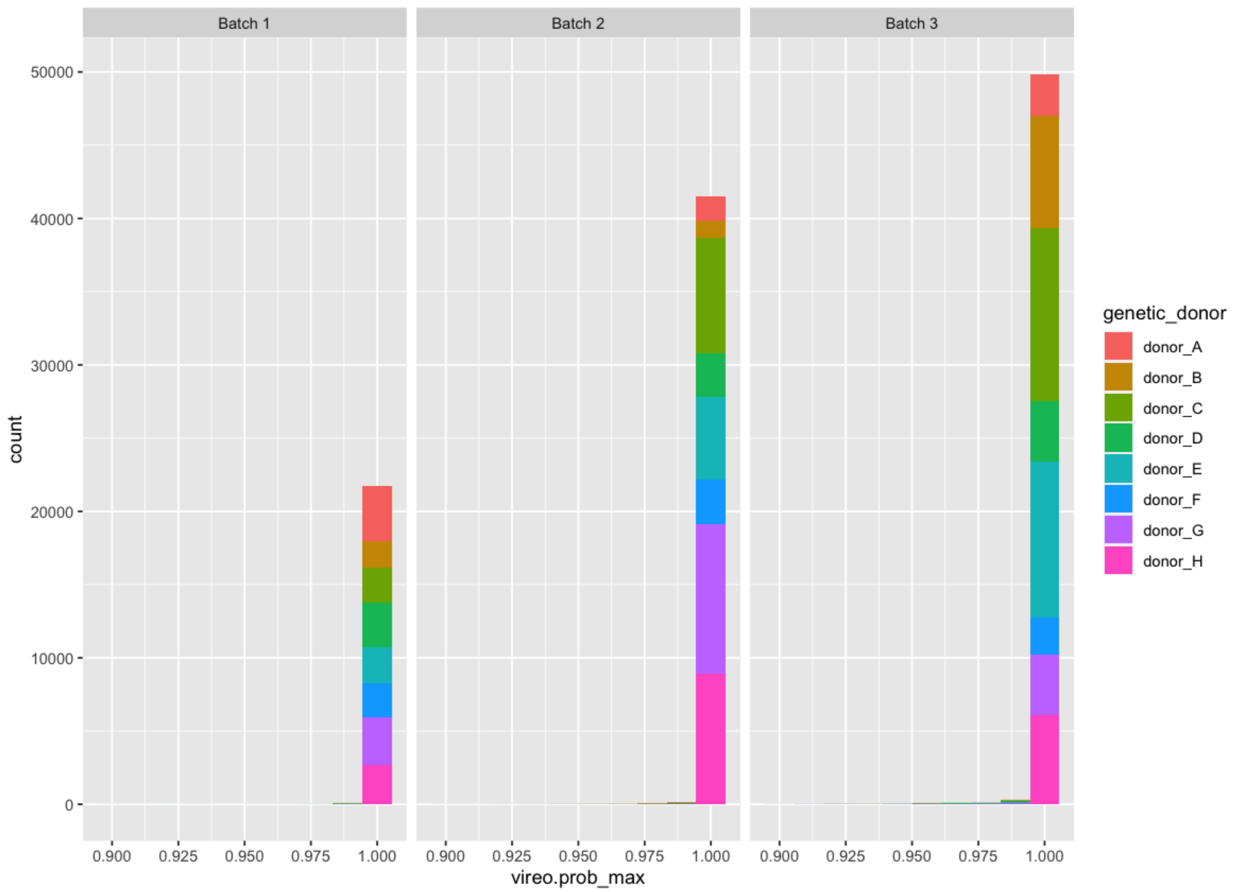


Figure S2: Singlet assignment confidence score from vireo for each batch in the first data set, coloured by donor.

Figure S2 shows that *vireo* performs well on this data set, giving a confidence score > 0.99 for 99.5% of the cells in all three batches. This is in contrast to the hashtag data, which is worse in batches 2 and 3 than in batch 1, as shown in Figure 1.

#### Parameter adjustment

Herein we describe three of the hashtag demultiplexing methods and determine how their performance on the BAL data set changes based on the value of certain tunable input parameters. As an overall performance metric, we compute only the median F-score for singlet assignment. As shown in Figures 4 and 5, the F-score is not a perfect metric for such complex data, and other measurements such as the fraction of true doublets erroneously assigned to singlet states are important to consider as well as the F-score. However, using multiple performance metrics necessitates deciding what weight should be given to each, something that will most likely change on a case-by-case basis. In the interest of simplicity, we use the F-score, as it provides a high-level view of the accuracy of assignment by each method and demonstrates that for most methods the default values are appropriate.

**hashedDrops**

The parameter that we find most affects the output of *hashedDrops* is `confident.min`. This is the value of the log-fold change between the most abundant and second-most abundant HTO for a single droplet, `LFC`. If `LFC` > `confident.min`, and it is not separately identified as a doublet, then the droplet is assigned as a singlet, and if `LFC` < `confident.min` it is assigned negative. The consequences of adjusting `confident.min` are fairly straightforward: higher values lead to greater precision at the cost of more false negatives, while lower values lead to more singlets at the cost of more false positives. Figure S3 shows the median F-score for singlets in batches 1, 2 and 3 of the BAL data set with 0 ≤ `confident.min` ≤ 3.

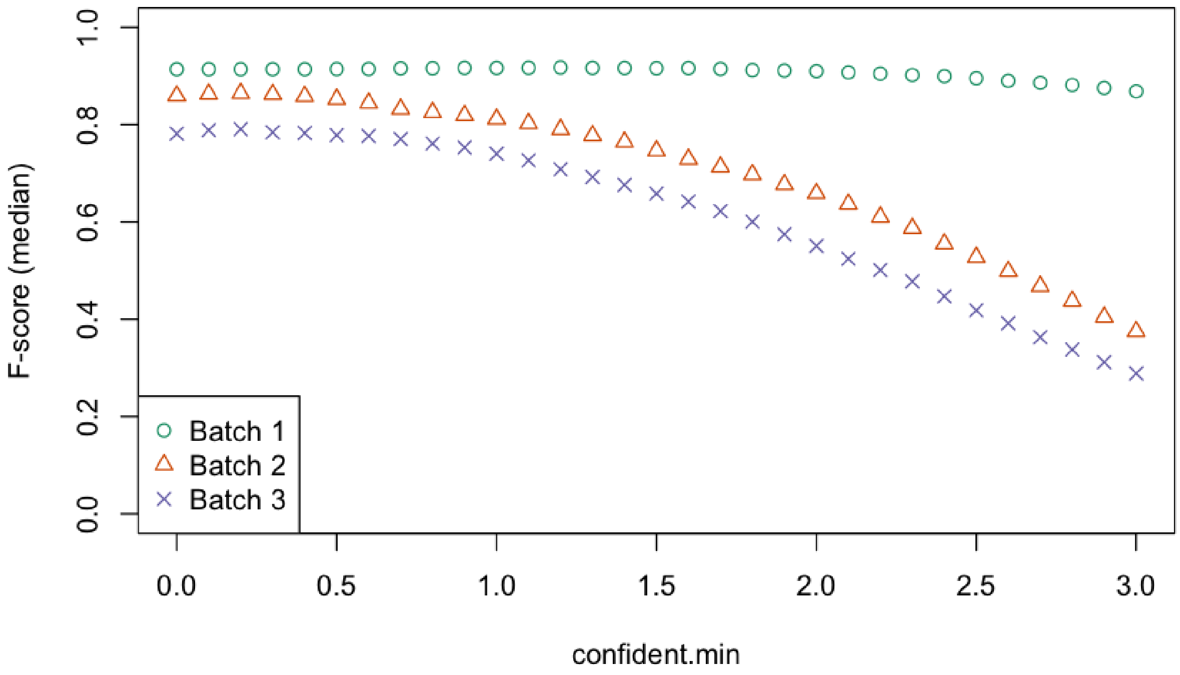


Figure S3: Median F-score from *hashedDrops* as a function of `confident.min` for batches 1 (green circles), batch 2 (orange triangles) and batch 3 (purple crosses) of the BAL data set.

Figure S3 shows that for batch 1, as shown in the results section and Figure 3, the F-score is insensitive to the choice of `confident.min`, while for batches 2 and 3, the F-score decreases for `confident.min` ≳ 0.5. For this reason, we consider both the default value of 2 and also 0.5 in the main analysis.

**HTODemux**

*HTODemux* operates by forming $N_{samples} + 1$ clusters, and for each HTO fits a negative binomial distribution to the cluster with the lowest value of counts for that HTO. The remaining droplets are compared to this distribution, and if the counts for a given HTO are above `positive.quantile` then that droplet is assigned as positive for that HTO. `positive.quantile` is the only tunable parameter in *HTODemux*. We consider 0.9 ≤ `positive.quantile` ≤ 0.99, and show the median F-score in each batch in Figure S4.

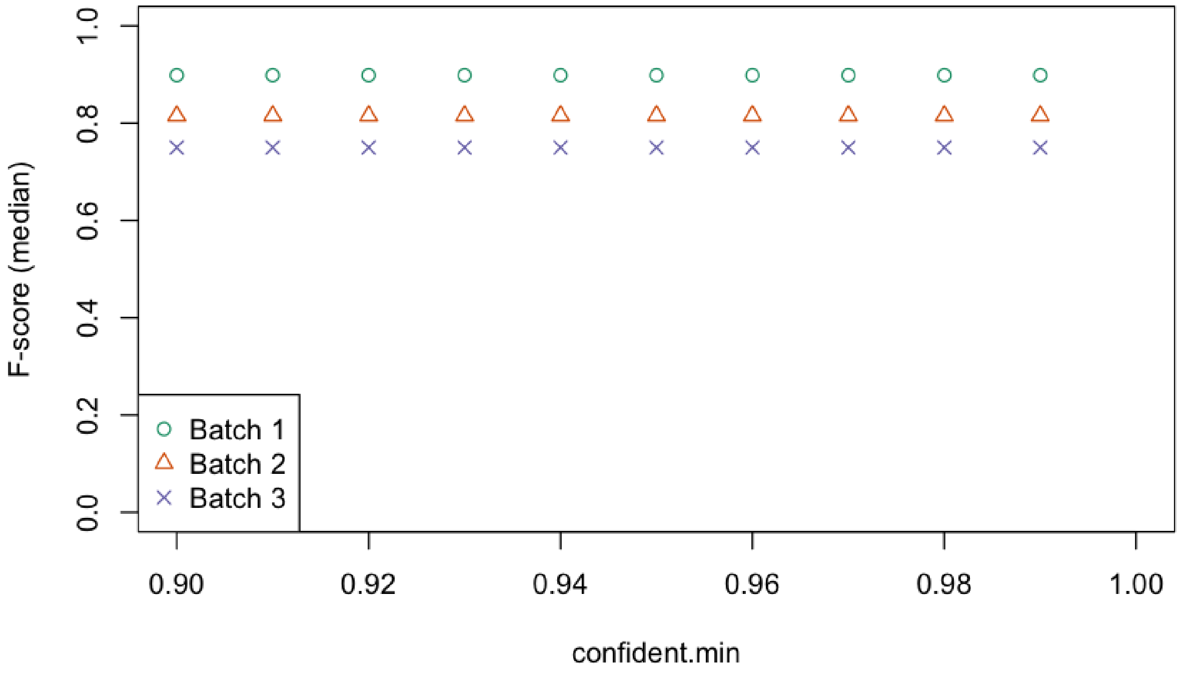


Figure S4: Median F-score from *HTODemux* as a function of `positive.quantile` for batches 1 (green circles), batch 2 (orange triangles) and batch 3 (purple crosses) of the BAL data set.

Figure S4 shows that the F-score is virtually insensitive to the value of `positive.quantile`, and so we use the default value of 0.99 in all of our analyses.

**GMM-Demux**

*GMM-Demux* produces a confidence score for each positive assignment of a droplet to an HTO. The threshold confidence score, *c*, that accepts these positive assignments is set to 0.8 by default, but can be adjusted by the user. We consider $0.5 \leq c \leq 0.99$ and show the median F-score for each batch in Figure S5.

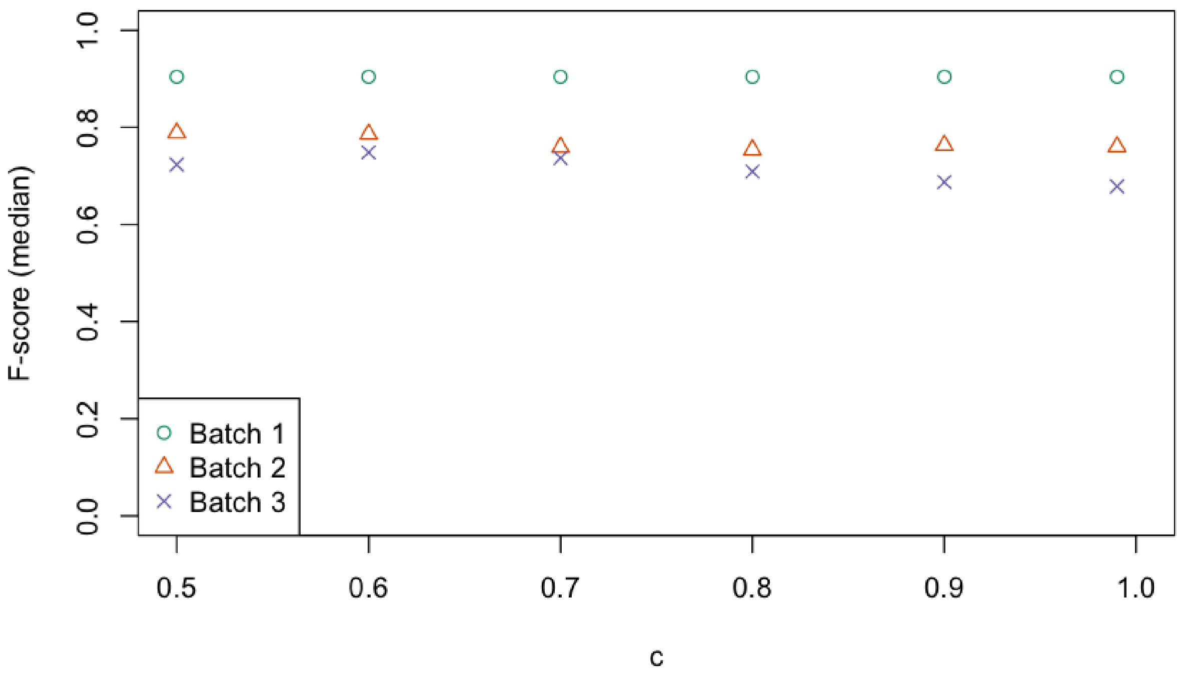


Figure S5: Median F-score from *GMM-Demux* as a function of minimum confidence threshold *c* for batches 1 (green circles), batch 2 (orange triangles) and batch 3 (purple crosses) of the BAL data set.

Figure S5 shows that the F-score is not significantly affected by the choice of *c*, and so we use the default value of 0.8 in all of our analyses.

**Ovarian tumour data**

In Figure S6, we show the PDF of HTO counts for each sample in each batch of the ovarian tumour data, as well as tSNE dimensionality reduction plots of the PCA of the normalized HTO counts, as in Figure 1 for the BAL data set.

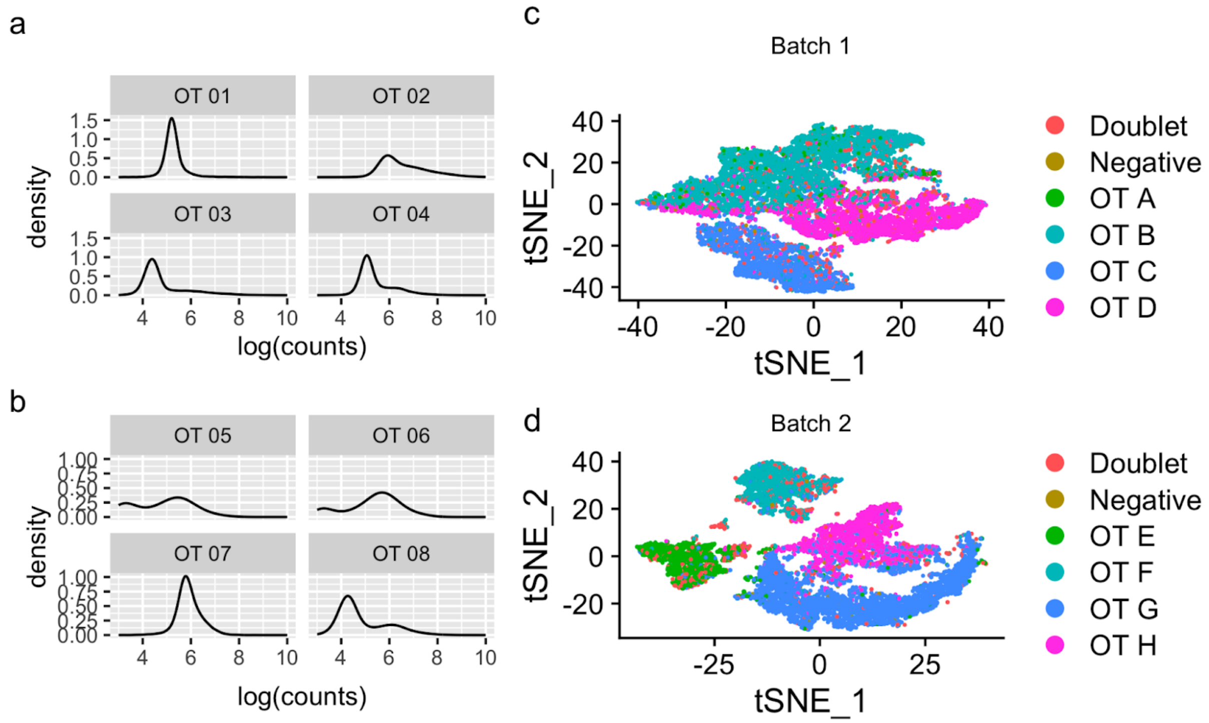


Figure S6: Quality assessment visualisations of the ovarian tumour data set. (a)-(b) PDFs of the logarithm of HTO counts for each hashtag, (a) is batch 1, (b) is batch 2. (c)-(d): tSNE dimensional reduction of HTO counts, coloured by genetic donor, (c) is batch 1, (d) is batch 2.

For some HTOs, in particular OT 01, OT 03 and OT 07, the second peak is not visible in the density plots (S6a and S6b). This indicates poor labelling of these samples, which is corroborated by the corresponding clusters in the tSNE plots (OT A, OT C adn OT G in Figure S6a and S6b) being poorly differentiated from the others, and from the low F-scores for these samples (Figure 5a in the main text).

**Cell line data QC visualisation**

In Figure S6, we show the PDF of LMO counts for each sample, as well as a tSNE dimensionality reduction plot of the PCA of the normalized LMO counts, as in Figure 1 for the BAL data set.

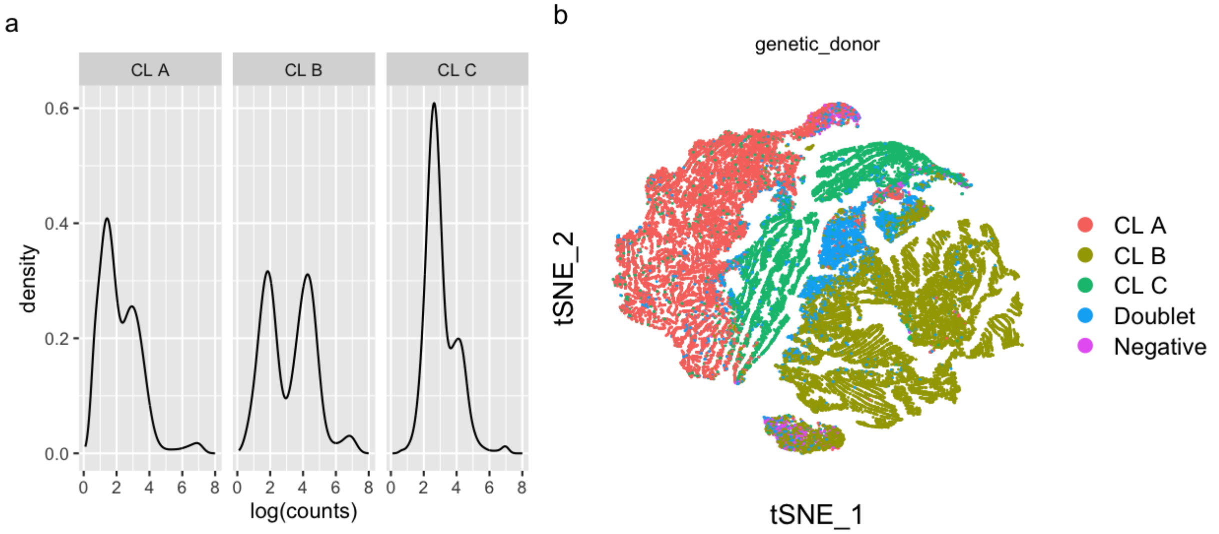


Figure S7: Quality assessment visualisations of the cell line data set. (a): PDF of the logarithm of HTO counts for each hashtag (labelled by corresponding genetic donor), (b): tSNE dimensional reduction of HTO counts, coloured by genetic donor.

As in Figure 1 (a), the PDFs in Figure S6 (a) show two peaks, corresponding to the background and positive distributions of each hashtag. Unlike in Figure 1 (a), however, the distributions in Figure S6 (a) show a third peak, likely corresponding to within-sample doublets. The size of this peak is proportional to 1 / $N_{HTOs}^2$, so in the BAL data set it is not resolvable above the noise. Figure S6 (b) is similar to Figure 1 (d-f), with three large clusters corresponding to each of the samples and the doublets and negatives interspersed throughout.
